# First detection of Crimean Congo Hemorrhagic Fever antibodies in cattle and wild fauna of southern continental France: investigation of explicative factors

**DOI:** 10.1101/2025.02.06.636810

**Authors:** Célia Bernard, Andrea Apolloni, Vladimir Grosbois, Armelle Peyraud, Phonsiri Saengram, Ferran Jori, Eva Faure, Nicolas Keck, Raphaëlle Pin, Olivier Ferraris, Loic Comtet, Benoit Combes, Matthieu Bastien, Valentin Chauvin, Laure Guerrini, Philippe Holzmuller, Laurence Vial

## Abstract

Crimean-Congo Haemorrhagic Fever (CCHF) is a tick-borne zoonosis with significant public health implications due to its expanding geographic distribution and potential for severe outcomes in humans. This study represents the first serological survey conducted in mainland France to detect antibodies against the Crimean-Congo Haemorrhagic Fever Virus (CCHFV) in domestic and wild fauna, providing critical insights into the virus’s circulation. We analyzed 8,609 cattle sera and 2,182 wildlife sera collected across the French Mediterranean region from 2008 to 2022, using enzyme-linked immunosorbent assays (ELISA) and pseudo-plaque reduction neutralization tests (PPRNT) for antibody detection and confirmation. Seropositivity was detected in both cattle (2.04%) and wildlife (2.25%), with higher rates observed in specific regions such as Pyrénées-Orientales and Hautes-Pyrénées. These findings reveal spatial clusters of CCHFV circulation and suggest the existence of enzootic transmission cycles involving local tick vectors and animal hosts. Our multivariate analysis identified key factors influencing seropositivity, including animal age, habitat characteristics, and potential wildlife interactions. The presence of natural open habitats and coniferous forests was significantly associated with higher seropositivity in cattle, while sex and geographical variability played a role in wildlife seroprevalence. These findings highlight the importance of environmental and anthropogenic factors in shaping the dynamics of CCHFV transmission. This study demonstrates that CCHFV is actively circulating in parts of mainland France, emphasizing the need for enhanced surveillance and integrated approaches to monitor zoonotic pathogens. It also raises questions about the role of additional tick vectors, such as *Hyalomma lusitanicum*, in the transmission cycle. These results contribute to a better understanding of CCHF epidemiology and offer valuable guidance for public health strategies to mitigate the risks associated with this emerging disease.

**Author Summary:** Crimean-Congo Haemorrhagic Fever (CCHF) is a severe disease spread by ticks that affects humans and animals. Although the disease is largely distributed in parts of Europe, Africa, and Asia, its presence in France has been uncertain. My study investigated whether the virus causing this disease is circulating in southern mainland France by testing blood samples from domestic animals, like cattle, and wild animals, such as deer and wild boars, for signs of previous infection.

I found evidence of the virus in several regions, particularly in the Pyrénées-Orientales, Hautes-Pyrénées and Alpes-maritimes, suggesting that the virus is indeed circulating among animals and ticks in some parts of mainland France. By studying where infected animals were found and considering factors such as age, habitat, and environmental conditions, I identified that older animals seem to have been more often exposed to the virus, as well as animals frequenting open environments favorable to ticks.

These findings are important because they show that the disease could potentially spread to new areas and affect human populations. My work highlights the need for ongoing monitoring of ticks and animals for CCHFV epidemiological surveillance and the protection of public health.

## Introduction

Crimean-Congo Haemorrhagic Fever (CCHF) is a tick-borne zoonosis, caused by the infection with a RNA virus of the *Orthonairovirus* genus and *Nairoviridae* family (1). CCHF is one of the most widespread tick-borne diseases with the widest distribution worldwide (2), which occurs in Africa, the Middle East, the Balkans, and Central Asia (3,4). Its recent detection into new areas such as India and Pakistan, as well as western Europe like Greece and Spain, over the last two decades is a public health concern (3,5,6). Humans are contaminated through tick bites, or when they are exposed to blood or infected tissues of viremic animals, or infected persons (WHO). Only part of infected people develop symptoms, with a first phase of fever, tremors, myalgia, headache, nausea and vomiting, abdominal pain and arthralgia, and sometimes complications with internal and external bleedings as well as ecchymosis; mortality of infected people varies between 9 to 50%, depending on the time of treatments implementation and the virulence of the viral strain (7,8). In contrast, the infection of Crimean-Congo Haemorrhagic Fever Virus (CCHFV) in wild or domestic animals does not cause any clinical signs, although most of infected species are able to become viremic and develop a humoral immune response (9). Although viremia in animals is short (5-10 days), antibodies against CCHFV are maintained for several years especially in large ungulates, making serology an excellent surveillance tool to measure animal exposure to CCHFV and detect early virus circulation in a naïve area (9,10). In addition, animals can play essential roles in CCHFV transmission, by either the amplification of tick vector populations, the transient replication of CCHFV to reinfect new tick vectors, or the spread of infected ticks (11,12). Animals can only be contaminated by infected ticks’ bites, and this circulation of the virus between vertebrate animals and ticks constitutes the enzootic natural transmission cycle of CCHFV (10). CCHFV genome has been detected in many tick species worldwide but only ticks of the genus *Hyalomma* (with *H. marginatum* and *H. lusitanicum* as likely vectors in southern Europe) are considered best competent vectors for CCHFV (13,14). Regarding their ability to remain infected during their development cycle, and their competence for horizontal but also transovarial transmission, as well as cofeeding, ticks are also considered as the natural reservoirs of CCHFV (15). Thus, the geographical distribution of CCHFV can expand through the introduction and establishment of infected ticks into new suitable territories, by either migratory birds or human-driven long-distance displacements of livestock. Since 2010, CCHF has been detected in western Europe, with virus genome detected in *Hyalomma spp.* ticks in Spain (16,17) and then the reporting of a few human cases linked to tick bites since 2016 (18). Antibodies against CCHFV have been detected in Spanish cattle and wildlife in Spain (19,20) but also in bovine sera from southern Italy (21). In France, a serological survey was conducted on domestic ruminants in Corsica, a French island southeast of the mainland, between 2014 and 2016, and detected about 9% of positive animals (cattle, sheep ang goat) among 4,000 tested (22). Given the recent establishment of the known tick vector *H. marginatum* on southern coastal areas of the mainland (22,23), the epidemiological situation of CCHF in this area is also questioned. Recently, CCHFV was detected for the first time in *H. marginatum* ticks in Pyrénées-Orientales on the mainland, as well as in Corsica Island, confirming the local transmission of CCHFV in France (24,25). Because the prevalence of tick infection is assumed to be very low in French tick vectors due to dilution effect (26), monitoring CCHFV in ticks remains laborious and needs large sampling effort, compared to serology, to determine virus circulation areas. In addition, knowing where are tick vectors is necessary but not sufficient to determine virus circulation locations, and other factors should be investigated. No studies have been carried out in Corsica to determine factors influencing seroprevalence, but research in that field has been carried out in other countries, and has identified a number of determinants impacting either the exposure of animals to infected tick vectors or the level of CCHV transmission within the enzootic cycle (27,28).

For this study, we focused on the French mainland where serological survey has never been carried out. Sera from cattle and wildlife were analyzed from different areas of the French Mediterranean rim, in order to detect antibodies against CCHFV and estimate individual seroprevalence in different animal populations as indicators of CCHFV circulation levels. Using a correlative approach, we also aimed to identify individual, environmental and anthropogenic determinants associated with CCHFV transmission.

## Results

### Seropositive animals using ELISA test

We obtained 8,609 cattle sera along four prophylaxis periods from 2018 to 2022 (Table 1), and 2,182 sera of wildlife from 2008 to 2022 (Table 2), to be tested using the ELISA test.

**Table 1:** Number of cattle, municipalities, and farms analyzed for detecting CCHFV antibodies, in each department that accepted to participate in the study, and number of cattle sera found positive in CCHFV antibodies.

| Department | Number of tested sera | Number of tested farms | Number of positive | Year of sampling |
| --- | --- | --- | --- | --- |
| Alpes-Maritimes | 571 | 75 | 41 (7 farms) | 2019-2020 |
| Aveyron | 1246 | 126 | 2 (2 farms) | 2019-2020<br>2020-2021 |
| Bouches-du-Rhône | 1710 | 188 | 7 (3 farms) | 2018-2019<br>2019-2020<br>2020-2021 |
| Gard | 1013 | 60 | 17 (2 farms) | 2018-2019<br>2019-2020 |
| Hérault | 1946 | 191 | 28 (10 farms) | 2018-2019<br>2019-2020 |
| Lozère | 100 | 20 | 0 | 2020-2021 |
| Pyrénées-Orientales | 880 | 87 | 80 (19 farms) | 2019-2020<br>2021-2022 |
| Tarn | 1112 | 111 | 1 (1 farm) | 2019-2020<br>2020-2021 |
| Other (Var and Vaucluse) | 31 | 4 | 0 | 2019-2020 |

**Table 2:** Number of wildlife sera obtained from 2008 to 2020, for each hunted animal species, analyzed for detecting CCHFV antibodies in each department who have collected sera on hunted animals and have accepted to participate in the study, and number of sera found positive in CCHFV antibodies.

| Department | Number of tested sera | Number of tested municipalities | Number of positive animals | Wildboar | Roe deer | Red deer | Mouflon | Fox | Year |
| --- | --- | --- | --- | --- | --- | --- | --- | --- | --- |
| Ardèche | 28 | 5/335 | 0 | 0/17 | 0/11 | - | - | - | 2009-2012 |
| Drôme | 434 | 35/363 | 0 | - | 0/434 | - | - | - | 2016-2020 |
| Gers | 379 | 98/461 | 0 | 0/74 | 0/305 | - | - | - | 2010-2021 |
| Hautes-Pyrénées | 621 | 148/469 | 45 | 9/179 | 16/197 | 20/244 | 0/1 | - | 2008-2022 |
| Hérault | 622 | 49/342 | 3 | 3/566 | 0/23 | 0/8 | 0/21 | 0/4 | 2012-2022 |
| Lozère | 98 | 12/152 | 1 | 0/33 | 0/7 | - | 1/58 | - | 2019-2022 |

Antibodies against CCHFV were detected by ELISA in 176 cattle sera in most of the tested departments, except in Lozère that remained negative (Table 1, Figure 2). Tarn and Aveyron only showed one and two seropositive animals, respectively, each of them isolated in a farm in separated municipalities. The two seropositive cattle in Aveyron are animals that have changed farms in the course of their lives: the first was born in Hérault in 2016 and arrived at the farm where it was sampled at the end of 2018. The second was born in Aveyron in 2011, and although it has changed farms, it has remained in the north-west of the department all its life. The seropositive animal sampled in Tarn has never changed farm since birth. Although individual seroprevalences were higher in Bouches-du-Rhône (0.4%), Hérault (1.44%) and Gard (1.68%), positive municipalities remained spatially dispersed and most of farms contained only one seropositive animal, except four farms that concentrated several seropositive cattle (intra-farm seroprevalence in farms with more than one positive animal 26.08% [4.08-66.67%]) (Figure 2). Then, individual seroprevalences were the highest in Alpes-Maritimes (7.18%) and Pyrénées-Orientales (9.09%), where seropositive municipalities were clustered around apparent “hotspots” of transmission (intra-municipality seroprevalence 39.1% [11.11-100%]) and most of positive farms concentrated many seropositive animals (intra-farm seroprevalences in clusters 47.93% [10-100%]) (Figure 2).

**Figure 1.**
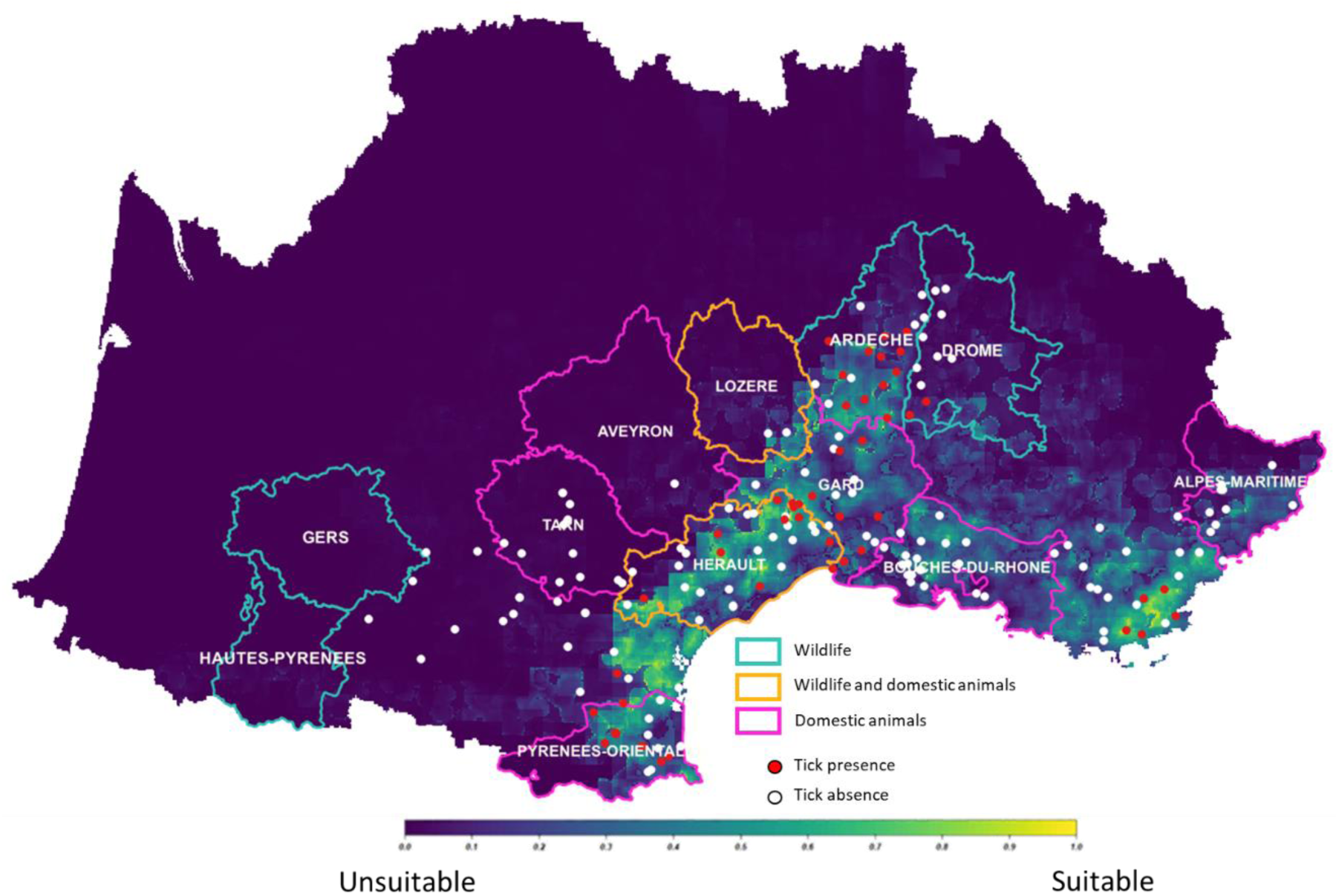
: Study area of the various participating departments in domestic and/or wild fauna. In the background, the map predicted probability of presence for *H. marginatum* in the South of France, using the most parsimonious model (dark blue: null probability to bright yellow: highest probability). Red circles represent the observed presences and white circles observed absences from testimonies and surveys. Adapted from (23).

**Figure 2:**
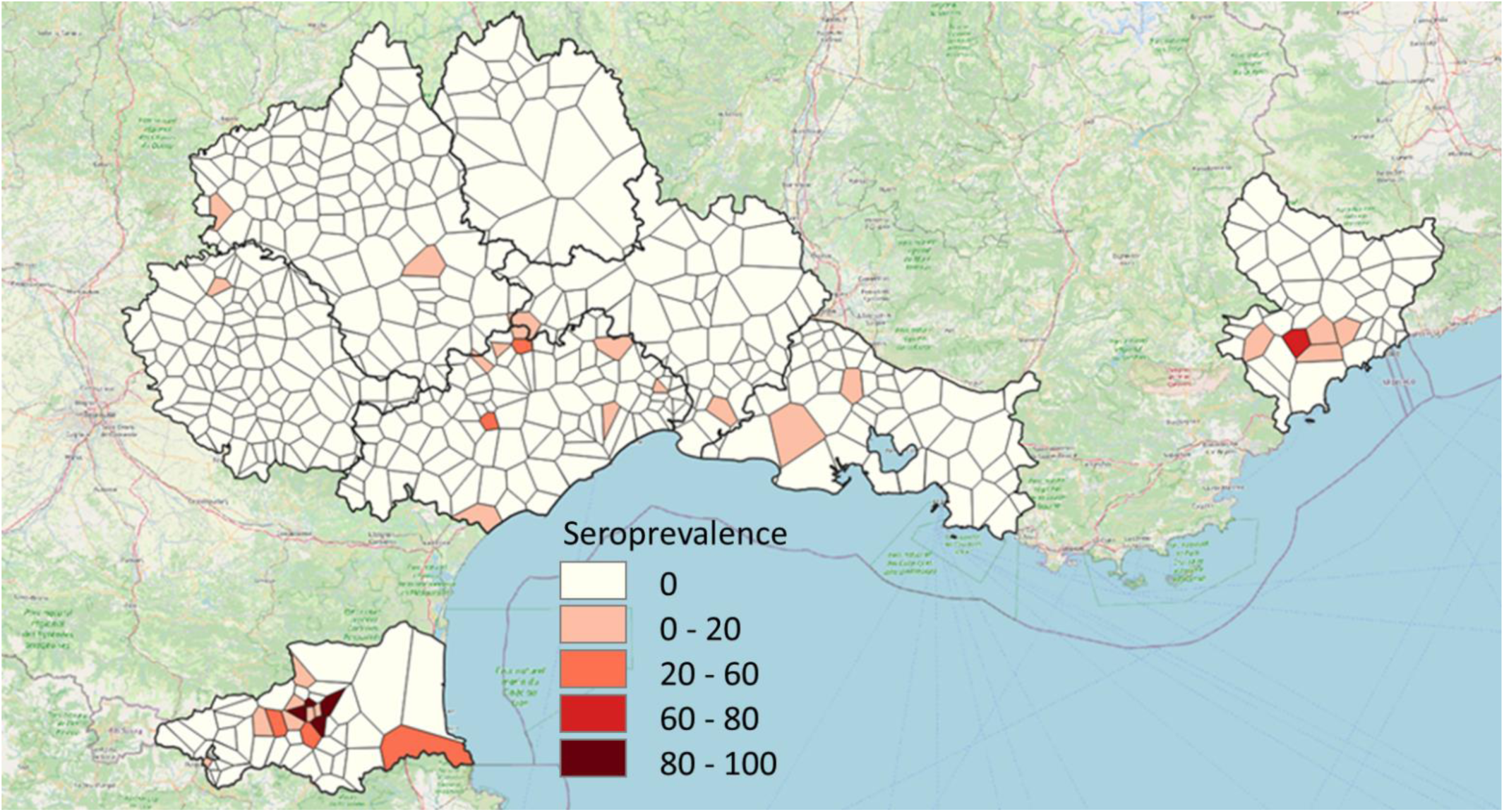
Map of cattle individual seroprevalences per municipality (increasing seroprevalences from pale pink to dark red, and sampled but negative municipalities in white), in departments that accepted to participate to the study. Voronoï polygons were used as a representation of municipalities, guaranteeing anonymity of farmers in sampled municipalities and dealing with the absence of sampling in some municipalities.

For wildlife, 49 positive samples were detected in wild boars (n=14), red deer (n=18), roe deer (n=13) and mouflon (n=1) (Figure 3). 45 of the 49 positive animals (including wild boar, roe deer and red deer) were hunted in Hautes-Pyrénées (Table 2). In this department, antibodies were detected almost every year since 2008, except during the COVID years (2019–2021) where hunting pressure drastically decreased.

**Figure 3:**
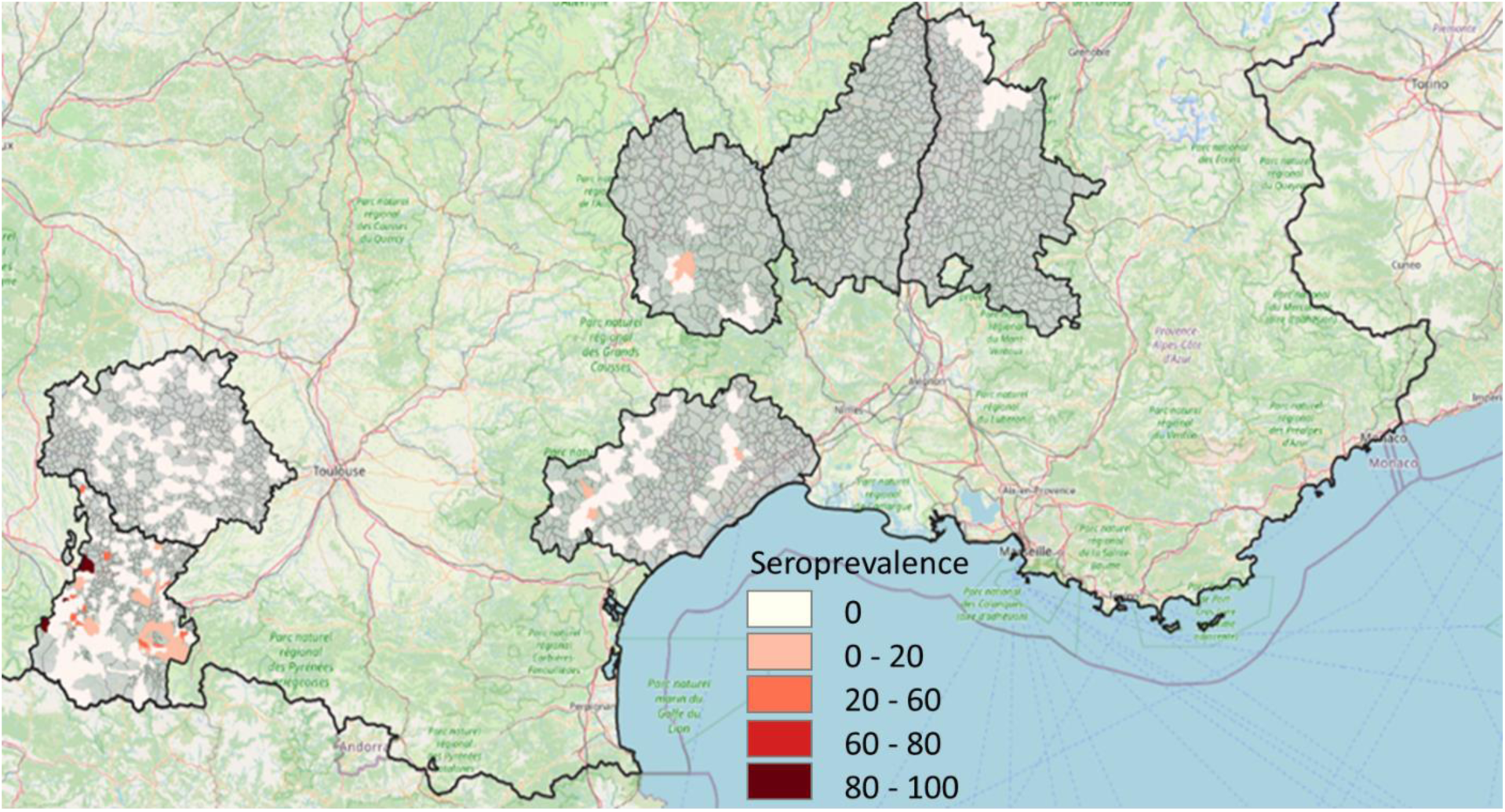
Map of wildlife seroprevalences per municipality in departments that accepted to participate to the study (increasing seroprevalences from pale pink to dark red, and sampled but negative municipalities in grey)

When looking at OD values of cattle sera tested by ELISA, positive and negative samples grouped in two well separated populations (Figure 4A). Negative samples exhibited a clustered distribution around a consistent OD value (median = 3.68). In contrast, positive samples exhibited two discernible profiles. Pyrénées-Orientales and Hérault demonstrated homogeneous high OD values (median = 152.36). Conversely, OD values for the other positive departments exhibited a broader distribution, with some samples approaching the ELISA threshold while others obtained higher OD values (median = 65.87). Among these samples, Alpes-Maritimes displayed a single prominent peak of OD values not so far from values obtained in Pyrénées-Orientales and Hérault, while Gard exhibited two distinct peaks. In the other departments, it was difficult to determinate profiles as positive samples were scarce but most of the OD values were low, close to the threshold. In wildlife, all negative samples exhibited similar low OD values (median = 5.93) (Figure 4B). As for cattle, positive samples displayed two distinct peaks of OD values, one peak aligning closely with the threshold and another with OD values (Figure 4B). This pattern was similar for the different animal species. The single mouflon found positive in Lozère had OD values close to the threshold.

**Figure 4:**
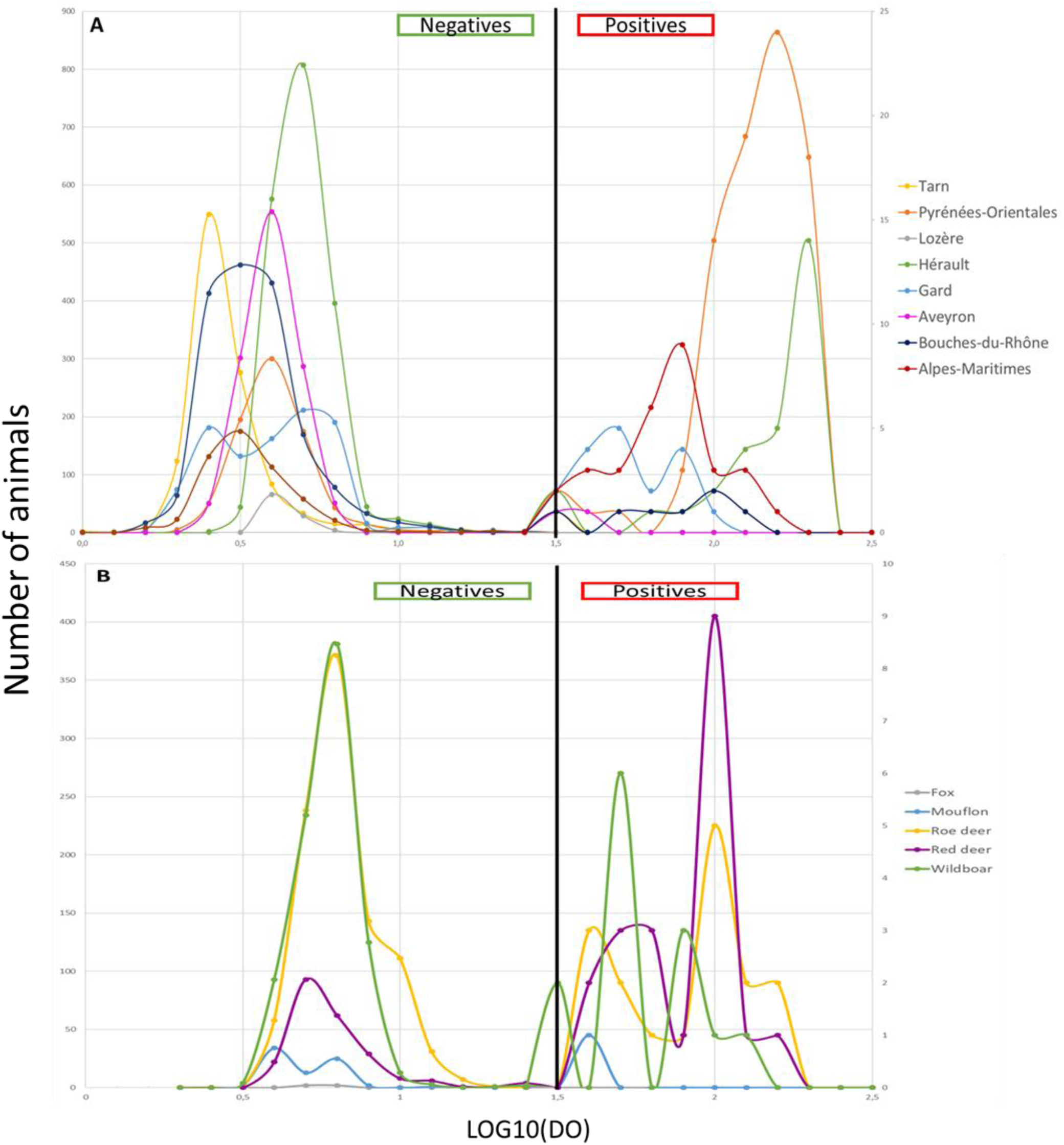
Estimated density distributions of positive and negative samples across different departments (A) and animal species (B), based on optical density measurements. The curves on the left represent negative samples, while those on the right correspond to positive samples, according to their optical density values. The curves illustrate the distribution profiles of optical density, highlighting differences across regions and species, and distinguishing positive from negative samples

### Confirmation of positivity using PPRN test

59 ELISA-positive cattle sera (Alpes-Maritimes = 17, Bouches-du-Rhône = 3, Gard = 10, Hérault = 20, Pyrénées-Orientales = 9) and 40 ELISA-positive wildlife sera (Hautes-Pyrénées = 39, Lozère = 1), in addition to 5 positive and 10 negative controls (as presented above), were sent to a BSL4 laboratory for conducting PPRN test. The results of PPRN are shown in supplementary data 1. Among controls, the 10 ELISA-negative samples were PPRN-negative, and 4 positive controls were confirmed positive by PPRN until 1/80 dilution and one at 1/20 dilution. Concerning cattle, 39 out of the 59 ELISA-positive sera were confirmed with a minimum threshold of 1/20. Those which could not be confirmed mainly came from Alpes-Maritimes (n=6/17), Bouches-du-Rhône (n=3/3), Gard (n=8/10) and a few from Hérault (n=3/20). Concerning wildlife, 39 out of the 40 ELISA-positive sera were confirmed with a minimum threshold of 1/20, and most of them (n=30/39) were still confirmed at 1/80 dilution.

There was a slight correlation between OD values provided by the ELISA test and Dilutions Thresholds (DT) provided by the PPRN test for cattle (R²=0.35) and wildlife (R²=0.29) (Figure 5A and 5B), suggesting that the higher the antibodies level is, the higher is the dilution until which it is possible to detect neutralizing antibodies. However, especially in cattle, some samples with high OD values remained negative in PPRN or were confirmed only at low dilutions.

**Figure 5:**
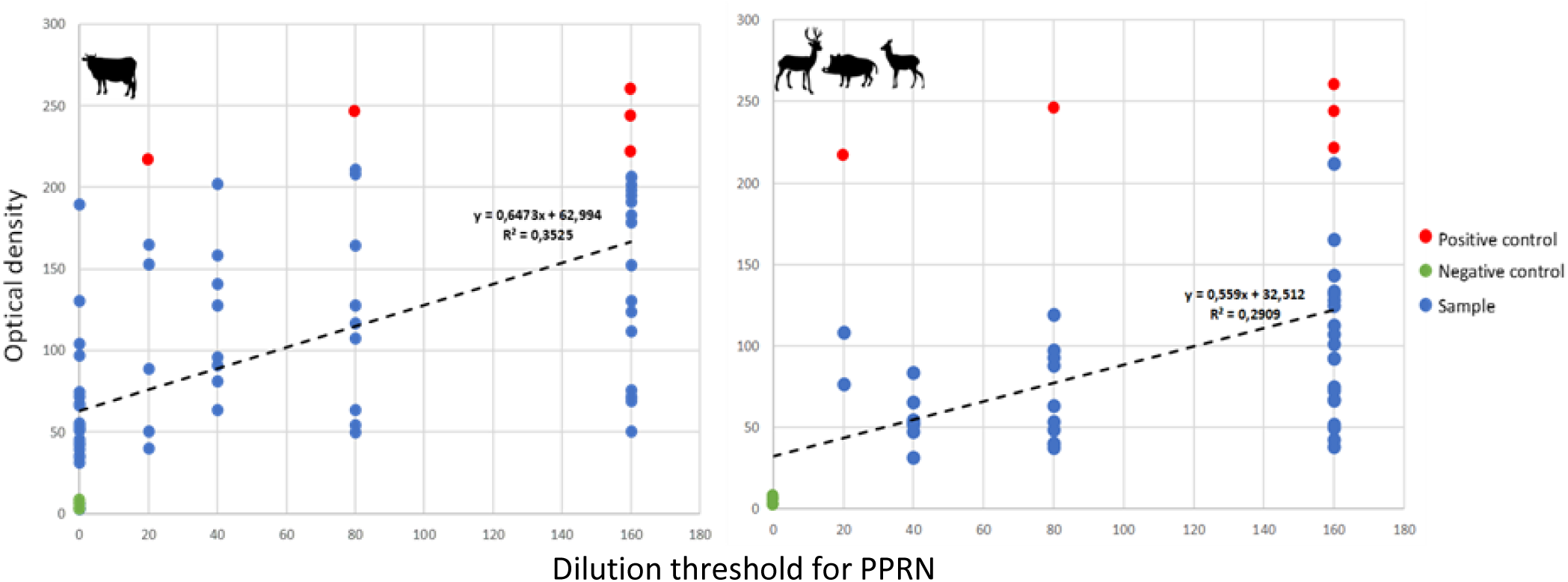
Correlation between the Optical Density (OD) of sera samples in ELISA and the Dilution Threshold (DT) until which the neutralizing effect of antibodies in sera is still observed by PPRN, in cattle (A) and wildlife (B). Reference positive controls (ELISA-positive Corsican sera) are indicated in red points and negative ones (ELISA-negative sera from Corsica and Netherlands) in green points.

### Factors explaining seropositivity in animals

In cattle, the effect of age was highly significant on animal seropositivity (P-value = 0.0004), indicating that as more as animals become older they are more likely to become infected and be seropositive. In addition, farms with an important proportion of open natural habitats showed significantly much more seropositive animals than locations where the other habitats were predominant (P-value = 0.027). The variables “sex” and “ Closed Coniferous Forest” were also retained in the final model but their effect were not significant, with males less seropositive than females and locations showing a predominance of closed coniferous forests with a higher proportion of seropositive cattle (Table 3). Concerning random effects, substantial variance was observed at the level of farm within municipality and department (variance = 47.473, std. dev. = 6.890), and in a less extent for municipality within department and department, suggesting significant spatial heterogeneity in serological status of farms even if they are close (Table 3). The variance explained by the breed of cattle is also low, indicating that some breeds of cattle are not likely to be infected through tick bite and seropositive. The final model has an AIC of 775.2 and a BIC of 838.7, indicating an acceptable fit of the data to the model. The model’s intercept is estimated at -12.619 (p < 2e-16), suggesting a very low baseline probability that an animal is seropositive.

**Table 3:**
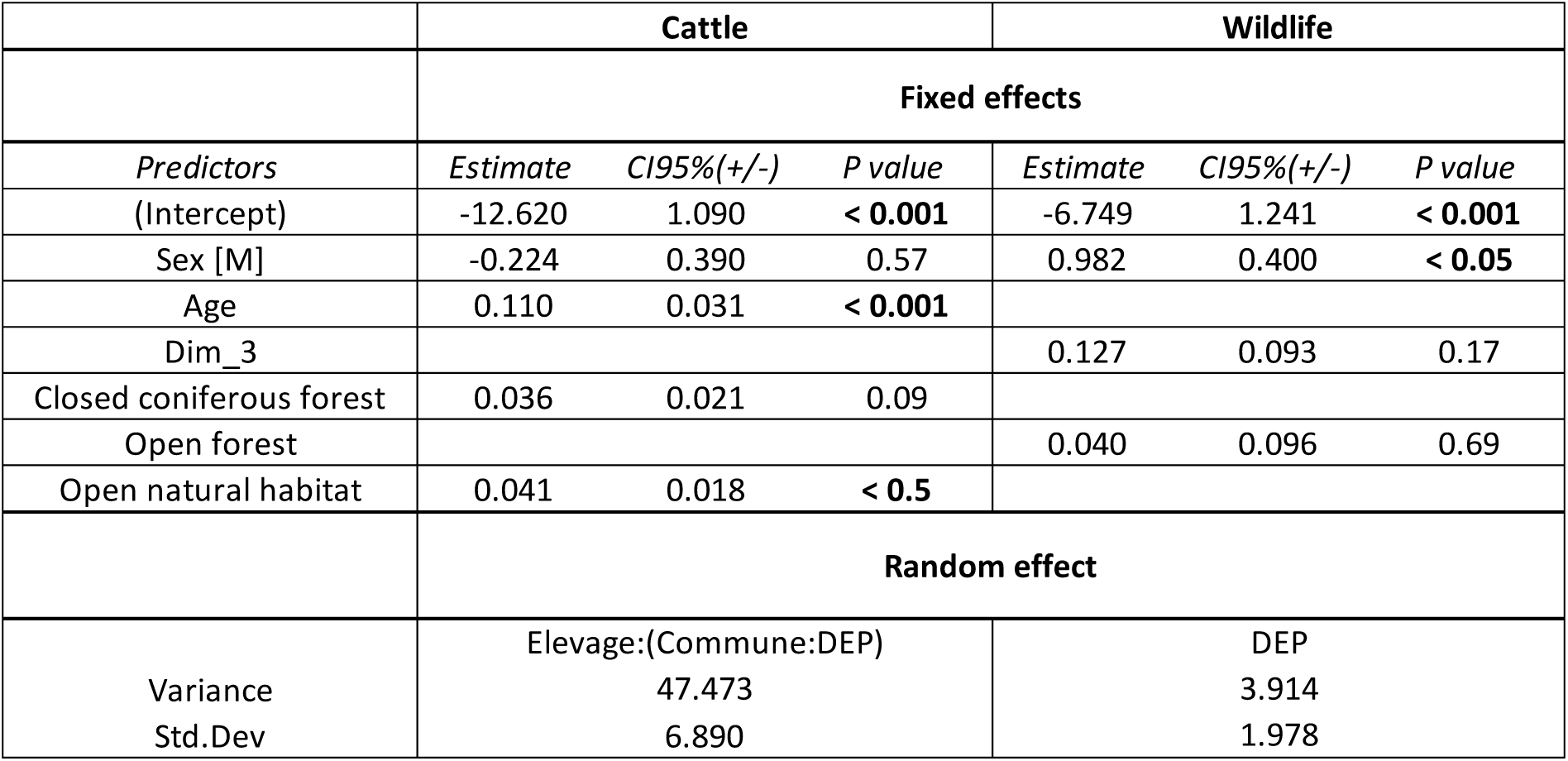
Factors retained in the final multivariate logistic regression model, for cattle and wildlife (according to models, rows are not filled in when variables were not retained), with the indication for each retained variable the tendency (estimate) and the significance of its effect (P-value)

In wildlife, less variables were retained in the final model and only the sex has a significant effect of the serological status of animals (*P*-value = 0.0141), with males being much more seropositive than females, at the inverse of cattle. The proportion of “Open_Forest” in municipalities was retained but showed no significant effect on seropositivity of animals (*P*-value = 0.6783. Similarly, the third synthetic climate variable “DIM 3”, mainly explained by autumn precipitation, was retained but did not show a significant effect on seropositivity (*P*-value = 0.1749). The random effects capture inter-departmental variability, with an estimated variance of 3.914. These results highlight the contribution of local factors and individual characteristics to the probability of infection, supporting the utility of mixed models in this context. The intercept of the model indicates both a baseline probability for a positive serological response and significant variations across geographical locations (departments). The simulated AIC for the model is 1523.45, and the BIC is 1540.78. These values indicate the trade-off between model fit and complexity, with lower values suggesting better model performance.

## Discussion

This work has led to the first detection of antibodies against CCHFV on mainland France, in cattle and wildlife, and allowed to determine areas of CCHFV circulation, at least in French departments where serological investigations were conducted. These novel results can question on the specificity of the ID Screen® CCHF DA ELISA we used in our study, which corresponds to the capacity of the test to only detect true positives. As mentioned before, by testing several sera samples from cattle, small ruminants and humans from CCHFV-free areas as well as sera from multiple animal species to confirm the absence of nonspecific reactions, a specificity of 100% (CI95%: 99.8%-100%) was determined (29). Despite this, one might wonder about potential cross-reactions with viruses genetically and antigenically close to CCHFV. A previous PPRN test carried out on ELISA-positive Corsican cattle sera has shown typical neutralizing effect of antibodies against CCHFV culture whereas no effect was observed with Hazara and Dugbe viruses, which belong to the same serogroup of CCHFV or a very close one (namely, the Nairobi Sheep Disease virus serogroup), respectively (Grech-Angelini, 2020). Some of these Corsican sera were included in our study and gave identical ELISA and PPRN results than the other tested sera from the mainland; discrepancy for some ELISA-positive sera that could not be confirmed by PPRN might result from the existence of different target epitopes between both tests (i.e. ELISA targets the nucleoprotein, while PPRN the glycoprotein) (30). In addition, another study conducted on cattle in Nigeria did not show any correlation between results provided by the ID Screen® CCHF DA ELISA and the DUGV ELISA developed to detect specific antibodies against Dugbe virus, while cross-reactions were reported when using immunofluorescence (31). Also, a survey where one calf and one sheep were hyper immunized with inactivated Hazara virus demonstrated that the ID Screen® CCHF DA ELISA was the only CCHFV serological test able to detect these animals as negative ones and thus to discriminate antibodies against Hazara virus and CCHFV (32). Similarly, a study showed that ruminants experimentally infected with Nairobi Sheep Disease virus were all negative for CCHFV antibodies when tested with the ID Screen® CCHF DAM ELISA (33). Regarding all these data, unless the existence of an unknown CCHF-like virus (sufficiently close to CCHFV to create cross-reaction in ELISA but sufficiently distinct to CCHFV to not be detected by CCHFV but also panNairovirus PCRs), our serological results suggest the circulation of CCHFV in the mainland France. With the recent detection of the causative virus in local *Hyalomma* ticks (24), the presence of persistent antibodies in ruminants confirmed the existence of an enzootic CCHFV transmission cycle between animals and ticks although human cases have never been reported yet in France.

In our study, different CCHF epidemiological patterns seem to occur among departments sampled in southern mainland France. In Tarn, Aveyron, Lozère and Bouches-du-Rhône, seroprevalences were very low, with maximum one seropositive animal per farm. OD values of positive animals were close to the threshold, and this could be interpreted as either false positive results, or single infections resulting in seroconversion that might be old compared to the date of antibody detection, with a possible decline of antibody titer, at least for the IgM that are also measured by the ELISA test used in this study. As the supposed CCHFV tick vector *H. marginatum* is predicted to be absent in these departments (23), they were thus considered CCHFV-free areas. Single infections could be due to human-driven displacements of domestic or wild ungulates from an endemic area where CCHFV already circulates, as it was already reported in other countries (26,34), or the introduction of an infected *H. marginatum* tick vector at the immature stage through bird migrations or introductions of lagomorphs for hunting and then molting of this tick in an adult that might be able to parasitize and infect a cattle locally (35–37). Punctual introductions of *H. marginatum* via trans-Mediterranean migratory birds used to be common in Europe and could explain most of single adult ticks found on horses at summer in northern countries, because of increasing temperatures allowing nymphs to molt in adult stages, without possibility for the tick to complete its entire development cycle and establish in these areas (38). In Gard and Hérault, seroprevalences and OD values remained low, seropositive municipalities were sparse, and the number of seropositive animals per farm was still close to 1, suggesting a similar non-circulating epidemiological situation. However, in some farms, several animals were seropositive with high OD values, which may suggest local CCHFV circulation with regular or recent infection of animals leading to persistent high virus titers. In these departments, *H. marginatum* is known to be present (22,23) and can explain such transmission. In Hérault, the fact of detecting a few seropositive wild boars, which are possible hosts for the adults of *H. marginatum*, also confirms the hypothesis of local foci of CCHFV circulation in domestic and wild ungulates. Finally, in Alpes-Maritimes and Pyrénées-Orientales, seroprevalences were significantly higher (7 and 9%, respectively) than in other departments and spatial clusters were identified around some municipalities, with many farms presenting a large number of positive animals. Such seroprevalences are quite similar to those observed in cattle from regions with supposed comparable epidemiological systems, like the French Corsica island (13.3%, 95%CI: 10.2%-17.3%) (39), in southern Italy (1.89%, 95%CI : 1.12–3.1) (21) or Central Macedonia in Greece (7%, 95%CI: 5%-10%). However, these seroprevalences remain very low compared to those observed in cattle in some CCHF-endemic African countries such as Mauritania (69%) or Mali (66%), where the epidemiological transmission cycles are different involving different tick vectors, with high parasitic loads on ruminants and various host preferences (40,41). Seroprevalences detected in wildlife of Hautes-Pyrénées (5.03% in wild boars and 8.16% in cervids) were slightly higher than those observed in same species on the other side of the Spanish border, in Catalonia (19), but they remain lower than those detected in central Spain where CCHF human cases have been regularly reported since 2016. However, in this last region, the epidemiological transmission cycle seems to likely involved the tick *H. lusitanicum* instead of (or in addition to) *H. marginatum* and consequently results are not totally comparable. In our study, OD values were medium to high in cattle from Alpes-Maritimes and Pyrénées-Orientales, as well as wildlife from Hautes-Pyrénées, with similar patterns than those observed in Corsican ruminants (39). All these results agree with frequent and active CCHFV transmission events between animal hosts and tick vectors in such departments of mainland France, although further field monitoring studies should be conducted to precise spatiotemporal dynamics of virus transmission. The seroprevalence in roe deer from this study, as high as in red deer, is the highest ever reported in the literature especially in South-Central Spain where this species has been found infested with infected *Hyalomma lusitanicum* (42). It is also worth mentioning that, despite CCHFV antibodies have been reported in other populations of mountain ungulates (19), this is the first time that mouflon (*Ovis musimon*) is detected seropositive in France, namely in Lozère. As it was a single individual, it remains uncertain if this was a false positive animal or just a single infection in such population after an *Hyalomma* tick introduction (see above).

Although *H. marginatum* is very abundant and largely distributed in Pyrénées-Orientales, in accordance with active CCHFV transmission in this department, such tick species has not been found in Alpes-Maritimes during two tick collection campaigns conducted on horses, respectively in 2017 (43). Similar discrepancy appears in Hautes-Pyrénées where seroprevalence in wildlife is high, whereas the tick *H. marginatum* is predicted to be absent in this department according to the distribution model developed by Bah et al. (23). Several hypotheses may be proposed to explain these situations. First, as the invasion of new French areas by *H. marginatum* in France is still in progress under climate changes, we may assume that these zones have become suitable for its establishment now, provided that it can spread from an endemic area and create sufficiently abundant populations to be detected by sampling. Another alternative would be the existence of another tick vector than *H. marginatum* in these departments. CCHF is a complex disease and, among its worldwide distribution range, its causative agent can be transmitted by various tick species belonging to the *Hyalomma* genus but also from other genera like *Rhipicephalus, Amblyomma,* and possibly *Dermacentor* (10,26). Recently reported as a likely vector of CCHFV in Spain (42), *H. lusitanicum* could be such a potential candidate for France. It was historically reported from France, in the western Atlantic region and Pyrénées-Orientales, which both surround Hautes-Pyrénées, and on the other side in Bouches-du-Rhône and Var near Alpes-Maritimes (44,45). It seems to be much more selective for its vertebrate animal hosts than *H. marginatum*, with a likely preference at the immature stages for lagomorphs, especially wild rabbits (46,47). As rabbit populations have greatly declined in the South of France since the 1970s with the epizooty of myxomatosis and more recently with rabbit hemorrhagic disease (36,37), we first assumed that this species may have disappeared from the territory (26). However, recent investigations detected a small population of *H. lusitanicum* in wild rabbit burrows in in Bouches-du-Rhône (Stachurski, pers. Comm.). This may be a residual population or a new population resulting from via rabbit introductions from Spain (37), and the extent of this tick species in France remains still unknown and needs further investigations in other French departments. If *H. lusitanicum* is in the future confirmed to be established in Hautes-Pyrénées, this might explain high CCHFV seroprevalences measured in roe- and red deer, which are likely hosts for the adult stages of *H. lusitanicum* (45,46). In Alpes-Maritimes, roe deer also exists in areas detected as CCHFV circulation hotspots, but further field investigations should be conducted both on tick distribution and serological survey in wild animals, in order to test such hypothesis. Another tick species that was frequently collected on horses in Alpes-Maritimes was *Rhipicephalus bursa*, which was proposed as a candidate vector for CCHFV although its vector competence has never been demonstrated (15). This tick species is also frequently encountered in sympatry with *H. marginatum* on horses and cattle in other southern French departments such as Pyrénées-Orientales, with similar seasonal activity at least for adult stages. In such region where CCHFV local transmission has been demonstrated, it would be interesting to investigate such tick species to assess its potential role in CCHFV transmission.

As *H. marginatum* is considered for instance as the only tick vector of CCHFV in France (24), the presence of CCHFV tick vector, namely *H. marginatum* for France (26), wasits presence was tested in our study to explain CCHFV seropositivity in cattle and wildlife, under the assumption that tick vector presence is one of the main environmental factors driving CCHFV natural enzootic transmission although tick vectors are not necessarily everywhere infected. By using predictions of *H. marginatum* presence from Bah *et al.* (23), our model did not detect any effect of tick presence . However, seropositivity in cattle was higher in areas with a high proportion of natural open habitats (i.e. shrublands in Mediterranean regions) and coniferous forests, and for wildlife, open forests was kept in the final model although the effect of this variable was not significant. Natural open habitats were already identified as most suitable conditions for the presence of *H. marginatum* (23), and may likely indicate optimal conditions for cattle of being infested by CCHFV tick vectors. However, other parameters than tick vector distribution may influence the efficiency of CCHFV transmission (26), such as the presence and the abundance of CCHFV-amplifying hosts, as well as spatiotemporal meeting occurrence between these hosts and the infected ticks. The fact that coniferous and open forests were pointed out by our two models might reflect the importance of wildlife, especially cervids, in CCHFV enzootic transmission cycle, as these animal species mainly colonize forested areas and were suspected to likely replicate CCHFV and reinfect new tick vectors in Spain (42). In addition, under the hypothesis of another tick vector like *H. lusitanicum*, and apart from the fact that abundant populations of cervids could contribute to amplify its populations, such forested areas may also provide direct suitable conditions for the survival and development of this tick species, as proposed by Valcarcel et al. (46). In this case, cattle in farms surrounded by such habitats would be likely to encounter infected tick vectors. Further research is therefore needed in these departments on the ecological niche of *H. marginatum* and other tick vector candidates, including also local interactions and likely associations between animals and tick species.

Apart from environmental conditions, anthropogenic factors related to farming practices were also tested to explain cattle seropositivity. Contrary to our assumption that the type of cattle farming (dairy, suckling, mixed, or leisure cattle) may influence the exposure of animals to tick vectors, this variable had no effect on cattle seropositivity in our study. In the South of France, most cattle are intended for meat production, and in peculiar areas such as Camargue some of them are also used for leisure activities, especially bullfighting (47). Consequently, we had an over-representation of suckling cattle compared to the other categories. In addition, it was sometimes complicated to attribute a breed to a category, especially for mixed uses. Finally, within the prophylaxis process as defined in France, dairy cattle are not blood sampled but milk is preferred to be tested for regulated diseases. As a consequence, this category was even more under-represented compared to suckling cattle in our study. Under this current study design, we cannot really conclude on such effect. Another anthropogenic factor that was tested in our model concerned contacts of cattle with other animal species, as these last ones may have different abilities for replicating CCHFV, contaminating tick vectors, and increasing CCHFV transmission in the vicinity of cattle (48,49). This variable showed no significant effect on cattle seropositivity, but it was not able to test contacts with wildlife as this information was not available from databases used in our study. This should be investigated through further field surveys by questioning directly farmers on their fencing practices or their observation of wildlife around farms. Such questionnaires to farmers could also inform on other practices such as their use of anti-parasitic products (insecticides, acaracides, or anthelmintics with ivermectine that is also efficient against ectoparasites such as ticks), a parameter that could not be included in our study due to lack of data. All these treatments are supposed to reduce tick infestation and thus decrease the exposure of cattle to infected tick vectors. This phenomenon was reported in cattle from Pakistan (49) and was considered a reliable preventive strategy when animals from Sudan were introduced into Saudi Arabia to avoid CCHFV transmission (34). Although no anthropogenic factors tested in our model showed an effect on cattle seropositivity, the fact that a large part of data variance was explained by the farm, as a random effect, confirmed that heterogeneity exists between farms, even within the same municipality, which could probably be explained either by the environment of farms or by farming practices. A last practice that is usual in the South of France, especially for suckling cattle, and should be further investigated is the temporary displacement of cattle towards spring and summer pastures during the activity period of the adult stages of *Hyalomma* ticks. As these pastures are usually far from farms and thus not described as the environment surrounding farms, this may be a limit of our current cattle model to identify habitats where cattle are more likely to become infested by CCHFV tick vectors. Studies conducted in Africa and Asia reported higher CCHFV seroprevalence in animals grazing in pastures than those fed on trough (48,49), and even higher prevalence in nomadic herds covering long distances (50).

Finally, individual factors were tested for explaining animal seropositivity in our survey. For both cattle and wildlife, the sex of animals was retained in final models, with females presenting higher seroprevalence than males in cattle, and the inverse for wildlife. Such observation is quite unusual and only two studies conducted on cattle from Malawi and South Africa reported higher seroprevalence in females than males (28,49). In our study, the over-representation of suckling cattle (see above) may have resulted in a bias favoring females than males, as reproductive females are usually kept longer than males that are sold for fattening or directly slaughtered after several months of life. Considering this bias, we cannot conclude on a reliable effect of sex on the probability for cattle to be seropositive. In wildlife, the larger home range of males than females (51), especially during the reproductive period, may multiply the opportunities for them to visit areas infested by CCHFV tick vectors. Another individual factor significantly affecting cattle seropositivity was the age of animals in our study. As already demonstrated in many serological surveys on CCHF in domestic ungulates, animals that are older have a higher probability to be seropositive, as a result of their longer exposure to infected tick vectors along successive years and the persistence of CCHFV antibodies for several years in most of ungulates (52). Conversely, age had no effect on wildlife seropositivity. However, in our study, it was categorized in only three classes and its assignment depends on the subjectivity of hunters, with possible difficulty to differentiate adults and subadults. Then, cattle breed as well as wildlife animal species were also tested as random and fixed effects, respectively, on the seropositivity. None of these variables had a significant effect and was retained in our best models. This is interesting as some studies conducted in Africa confirmed the apparent resistance of some cattle breeds, especially native ones, against tick infestation, and thus CCHFV transmission (53), but this tendency was not systematically highlighted (27), and in some cases just the opposite (48). In France, considering that the assumed tick vector *H. marginatum* has recently colonized the territory, it seems that no local breed had time to develop adaptive resistance to tick infestation and this is in agreement with our results. Regarding wildlife, the absence of effect for animal species seems to suggest that wild boar, roe deer and red deer were equally infested by CCHFV tick vectors, at least in Hautes-Pyrénées where most of seropositive animals were detected. This reinforces the importance of further surveys on tick vectors’ distribution and trophic preferences, as well as large serological investigations on the different wildlife species in the other departments.

### Ethic statement

Our study was strengthened by the willing participation of farmers, who consented to the analysis of sera collected from their livestock for the detection of CCHFV antibodies. In most of departments, they were asked individually, citing confidentiality and the absence of consequences for seropositive farms. Only in Tarn, the GDS (Groupement de Défense Sanitaire) that is the departmental institution representing farmers took the responsibility of the analysis. To ensure the privacy and confidentiality of data, robust data protection measures were implemented during sample handling, storage, and analysis. Ethical considerations extend to the use and conservation of these precious samples, in accordance with guidelines established with CIRAD’s legal department..

## Materials and methods

### Study area/period and sample selection

The study area was focused on South-Eastern France, from Spain to Italy, in areas with a confirmed presence of *H. marginatum,* or close areas predicted as suitable for its potential establishment, in the more or less long term, due to climate changes (22,23) (Figure 1). Some neighboring localities where *H. marginatum* was a priori absent were also selected, as the establishment process of this exotic tick species is still ongoing (23). Sampling was carried out at the department (corresponding to NUTS-3 administrative division in the European nomenclature) and municipalities (lower division within departments) levels. Most of these departments home a large number of cattle, particularly suckler cows for meat marketing. There are also a number of local breeds, such as fighting bulls and cows in Camargue, a humid area in Delta of Rhône. Wild boar is predominant all over the study area, while cervids such as roe and red deer become numerous when leaving the littoral, in dense forests from the hinterland.

Sera from cattle were selected among those collected during the national prophylaxis program. Indeed, as part of the national surveillance of regulated animal diseases, domestic livestock (large and small ruminants) are blood sampled every winter to be serologically tested for several pathogens. Blood samples are collected by official designated veterinarians in each French department, and sera are analyzed by official veterinary departmental laboratories. For our study, we thus needed the agreement of laboratories to participate and deliver samples after their use, and selection of samples first depended on their own sample management and storage policy. Considering this primary constraint, a subsampling of sera was conducted, in order to obtain large spatial covering in concerned departments and be able to detect positive farms and estimate animal seroprevalence, at least at the municipality level As *H. marginatum* is distributed in spatial clusters and seems to continue spreading (23), we predicted small local foci of CCHFV circulation if existing, with no assumption on the level of spatial heterogeneity. For this reason, in participating departments, all municipalities with cattle farms were included when possible and all farms in these municipalities (or at least 6 farms per municipality) were tested. In the previous serological investigation in Corsica, more than 25% of farms had at least one positive cattle (Grech-Angelini, personal communication) but we expected a much lower proportion of seropositive farms on the mainland due to the recent establishment of *H. marginatum*. In each farm, the number of sera was limited to 5-10 animals, which was considered enough to conclude on the epidemiological status of farms, as the estimated individual seroprevalence in positive farms was higher than 30% in Corsica (Grech-Angelini, personal communication). Sera were selected from three successive prophylaxis campaigns (2018-2019, 2019-2020, and 2020-2021) (Table 1) to cover as many farms as possible, according to available samples in laboratories. Only for Pyrénées-Orientales, a second 2021-2022 prophylaxis period was added to complete our sera sampling. In our analysis, because of the long persistence of CCHFV antibodies in most vertebrate animals (9), and the fact that prophylaxis campaigns were different from one department to another, we considered all these successive years as the same single period in order to allow comparisons. In the case of the few animals that were sampled twice (N=262), we only considered their last serological status, as no animal became seronegative from one year to the other and only 3 animals seroconverted.

Sera from wildlife were facilitated by the national hunter’s federation from some departmental hunting associations that accepted to participate in the study (Figure 1). Sera collections were available from several hunting seasons, from 2008 to 2022, and included samples from wild boar (*Sus scrofa*), mouflon (*Ovis orientalis*), red deer (*Cervus elaphus*), roe deer (*Capreolus capreolus*), and fox (*Vulpes vulpes)* (Table2).

### Serological analysis of samples

#### Enzyme-Linked Immunosorbent Assay (ELISA)

All sera selected from cattle and wildlife were tested for the presence of IgG and IgM antibodies against CCHFV, using a double-antigen ELISA kit (ID Screen CCHF Double Antigen Multispecies, Innovative Diagnostics (formerly IdVet, https://www.innovative-diagnostics.com) according to the manufacturer’s instructions. For this kit, the 95% CI for sensitivity is 96.8%–99.8%, and 95% CI for specificity is 99.8%–100% (29). A freeze-dried serum provided by the manufacturer (MRI-CCHF) was used as an external control to ensure that analytical sensitivity remained constant between runs and to determine the test uncertainty. Briefly, after reconstitution, the serum was serially diluted to obtain the detection threshold. Aliquots (3 freezing/thawing cycles max. per aliquot) were prepared and analyzed on each test run. For each serum sample, the absorbance of the colorimetric immuno-enzymatic reaction of the sample was divided by the absorbance of the positive control to obtain a percentage as the Optical Density (OD) of the sample. Samples with OD < 30% were considered negative and those with OD ≥ 30% were considered positive.

### Pseudo-Plaque Reduction Neutralization (PPRN)

In order to confirm serological results obtained by the ELISA test, we sent our ELISA-positive sera as well as 10 ELISA-negative sera (5 from Corsica and 5 from the CCHFV-free Netherlands) to a Biosafety Level 4 laboratory (Laboratory Jean Mérieux, Lyon, France) to be analyzed by the national reference laboratory for CCHFV (Institut de Recherche Biomédicale des Armées, Paris, France) designated by the World Organisation for Animal Health. They used the Pseudo–Plaque Reduction Neutralization test (PPRNT) (6) to measure the neutralizing effect of sera antibodies against the IbAr10200 CCHFV strain (same antigen used in the ELISA test), at successive dilutions 1/20, 1/40, 1/80, 1/160. Results were given by Dilution Threshold (DT) until which the neutralizing effect of each sera sample was observed. We also included in our confirmation process 5 ELISA-positive cattle Corsican sera for which the absence of immune cross-reaction with CCHFV close-related viruses (namely, the Hazara virus from the same serogroup as CCHFV and the Dugbe virus that belongs to the close Nairobi sheep disease serogroup) was already confirmed in previous CCHF survey, although they clearly neutralized CCHFV culture (39).

#### Statistical analysis of serological data

Considering the importance for protecting personal data of farmers and the risk of demonizing some farms, it was not possible to represent serological results in cattle by farms. In addition, taking into account the variability of sera samples’ availability across localities, it was biased to present data by municipalities and we preferred using Voronoi polygons that allow obscuring administrative divisions and to delineate new regional boundaries in accordance to sera sampling effort. This innovative approach facilitated the estimation of seroprevalence distribution and was conducted using the Voronoi function in QGIS. This technique involved generating Voronoi polygons, which assign regions based on proximity to sampled points, effectively avoiding the influence of administrative boundaries. The QGIS tool provided the flexibility to create these polygons efficiently, ensuring a more unbiased visualization of the data distribution across the study area.

In order to identify plausible factors linked to animal seropositivity against CCHFV in France (previously cited in (26)), we modeled the individual status of animals against their own characteristics, as well as environmental and anthropic factors. Individual information and rearing practices were extracted from the National Identification Database (BDNI) for cattle, and individual characteristics from hunting datasheets for wildlife, while environmental data were downloaded from previously compiled data by Bah et al. (23).

Concerning individual characteristics, for cattle, we informed their sex (male or female), age at blood collection (in years) and breed. For wildlife, we informed their sex (male or female), age (estimated as young, subadult or adult) as well as the species. The year when animals were sacrificed was also mentioned, as sera analyzed in this study have been collected for 14 years, and CCHFV transmission as well as host dynamics may have changed among this period.

Concerning rearing practices, for each farm of origin of tested cattle, we informed the rearing of other domestic animals (i.e. sheep, goat, pigs, or a mix of these) in addition to cattle on the farm, as an index of contacts for cattle with different animal species that may have distinct abilities to replicate CCHFV and infect tick vectors. Cattle breeds were also grouped to create categories that reflects the different types of production, for which we assume distinct grazing practices and various exposure to CCHFV tick vectors (dairy cattle mostly in barns, rustic dairy cattle mixing barns and pastures, free-ranging suckling cattle for meat production, peculiar cattle breeds used for leisure such as bullfighting or demonstrations).Concerning environmental data, the probability of *H. marginatum* presence in the South of France, derived from the suitability of local habitats and climate for the establishment of this tick species (23), was the first information to be collected, as an index of the exposure of animals to CCHFV tick vectors. However, we also considered the alternative scenario where *H. marginatum* would not be the only or the main vector of CCHFV in France, as it seems to be the case in Spain (54), and we thus also collected raw data on climate and habitat as index of tick vector presence. Indeed, each tick species has its own environmental needs to survive and succeed its development cycle, which defines its spatio-temporal ecological niche (38). In addition, climate and/or habitats may impact not only the dynamics of tick vectors but also other parameters of CCHFV transmission such as the distribution, the abundance or the specific diversity of vertebrate animal hosts, as well as the success of virus replication in ectothermic tick vectors (26). For climate, we used synthetic variables already defined by principal component analysis, which processed meteorological data in the South of France, from 2000 to 2018, and including temperatures, rainfall, potential evapotranspiration and relative humidity (23): Dim 1 (positively correlated with lower temperatures all the year and higher summer rainfall), Dim 2 (negatively correlated with high relative humidity and winter rainfall, positively correlated with evapotranspiration), Dim 3 (positively correlated with autumn rainfall). For habitats, we used a raster file (in pixels) presenting 7 land cover classes (humid areas such as inland marshes and rice fields, open natural areas such as sclerophyllous vegetation and sparsely vegetated areas, open forests, broad-leaved forests, coniferous forests, urban or peri-urban areas, agricultural areas) generated by Bah et al. (23), through the combination of CORINE Land Cover (CORINE Land Cover 2018, Version 2020_20u1) and BD forêt (BD Forêt® version 2.0). To attribute climate and habitat, as well as the predicted probability of presence for *H. marginatum*, to each farm from which cattle were serologically tested, we preferred considering the potential area where they could graze instead of the local point of the farm. Using the GPS coordinates of the farm’s head office, we created a 4-km radius buffer zone around each GPS coordinate. Within this buffer, we calculated the average *H. marginatum’*s probability of presence, and mean values for the three climatic principal components. For the habitat, we calculated a proportion for each of the 7 classes in each buffer zone. As data from wildlife were located at the municipality level, the similar measures were calculated for climate, habitat and *H. marginatum*’s probability of presence for each municipality where wild animals were serologically tested. The QGIS software (3.28.3-Firenze) was used for processing spatial dataset (“Zonal statistics” and” Zonal Histogram”).

To estimate the effect of these factors on the probability for an individual animal to be seropositive, we developed a Hierarchical Multivariate Logistic Regression Generalized Mixed Effect Model, respectively for cattle and wildlife, with a logit link function. The first concerned cattle with several nested levels (department, municipality within department and farm within municipality) and the breed (30 reported breeds) as random effects, and the other explanatory variables (sex, age, possible contacts with other animal species on farm, rearing practices concerning pasturing, *H. marginatum*’s probability of presence, climate, and habitat) as fixed effects. The second model concerned wildlife, with two levels of random effects (department and municipality within department), and the other explanatory variables (sex, age, year of hunting, *H. marginatum*’s probability of presence, climate, and habitat) as fixed effects.

Model processing was performed using R programming language (Rstudio, version 4.2.1). We used a dredge procedure (MuMin package) adding individual, anthropogenic and environmental fixed factors to select the most parsimonious model and only retain important explanatory variables. The optimal model was chosen using a combination of evaluation criteria: the AIC (Akaike’s information criteria) to gauge the overall quality of the model, the percentage of explained variance of data by the model, the Somers Dxy rank correlation test (Packages: car, MuMin, Lme4) to evaluate predictive capability of the model, and the AUC (Area under the ROC Curve) to assess its explanatory capacity. Taking into consideration these various measures, the model that demonstrated the best balance between data fit, predictive ability, and explanatory power was selected as the top choice.

## Acknowledgment

The authors would like to thank participating departmental veterinary laboratories and hunting departmental federations for their help to provide samples. We would also like to thank the management of health protection groups in the departments, to obtain breeders’ agreements. We would like to thank all cattle owners for collaborating with us on this study. We would like to thank the French Ministry of Agriculture to give us the access to the national database for livestock identification. Finally, we would like to thank the support services in CIRAD to help us in the process of protecting cattle owners individual informations.

